# Mother knows best: maternal egg-clutch size predicts larval group size in pine sawflies (genus *Neodiprion*)

**DOI:** 10.1101/761528

**Authors:** John W. Terbot, Catherine R. Linnen

## Abstract

Evolutionary conflicts are pervasive in nature and have the potential to drive antagonistic coevolution of conflict-related traits. However, when such conflicts are weak or idiosyncratic, phenotypic signatures of coevolutionary arms races may be absent. Here, we ask whether variation in group-living traits among pine-sawfly species in the genus *Neodiprion* is consistent with a history of parent-offspring conflict. To address this question, we compile data on adult female clutch size, larval aggregation behavior, and larval group size for a monophyletic group of 19 eastern North American *Neodiprion* species from field observations, laboratory assays, and published descriptions. We then evaluate the extent to which each trait exhibits phylogenetic signal and, based on these results, examine correlations between group-size traits both with and without phylogenetic correction. Although female oviposition behavior and larval grouping behavior varies among species and variation in these traits is decoupled from phylogeny, we find no evidence of antagonistic coevolution between these traits. Furthermore, while larvae are physically capable of dispersal, female clutch size is a strong predictor of larval colony size, indicating that larvae do not substantially alter initial group size after hatching. Thus, although theoretical work demonstrates the potential for parent-offspring conflict over group size in animals that lack parental care, our data suggest that this type of conflict is not likely to be a long-term driver of phenotypic evolution.

## Introduction

Evolutionary theory predicts that conflict—scenarios in which interacting entities have divergent fitness optima for some shared trait of interest—is widespread in nature (Queller and Strassmann 2008). Conflict manifests in a variety of contexts, including between males and females (sexual conflict; (Chapman et al. 2003; Arnqvist and Rowe 2005), parents and their offspring (parent-offspring conflict; (Trivers 1974; Godfray 1995; Crespi and Semeniuk 2004)), and within the genome of a single individual (intragenomic conflict; (Burt and Trivers 2009; Garder and Úbeda 2017)). Regardless of the context, evolutionary conflicts have the potential to fuel antagonistic coevolution, in which there is reciprocal evolutionary change in the genes and traits that mediate the conflict (Rice 1996; O’Neill et al. 2007; Carmona et al. 2015; Lindholm et al. 2016; Dougherty et al. 2017). When this coevolutionary arms race plays out over the course of lineage diversification, it can give rise to correlated evolution of conflict-related traits (Briskie et al. 1994; Arnqvist and Rowe 2002; Lloyd and Martin 2003; Koene and Schulenburg 2005; Kölliker et al. 2005; Ronn et al. 2007) and elevated rates of diversification (Arnqvist et al. 2000; Zen and Zen 2000; Kraaijeveld et al. 2011; Crespi and Nosil 2013). Thus, a comparative approach is a potentially powerful tool for making inferences about the strength and long-term evolutionary consequences of conflict.

One type of conflict that has received theoretical attention, but minimal comparative study, is conflict between parents and offspring over offspring group size. Whenever siblings compete for limited resources, a female’s optimal clutch size can differ from that of her offspring (Godfray 1987; Parker and Mock 1987; Godfray and Ives 1988; Damman 1991; Godfray et al. 1991; Desouhand et al. 2000). This type of parent-offspring conflict could occur even in the absence of any parental care (Costa 2018). For example, siblicide among parasitoid larvae may constrain females to produce clutches smaller than would be otherwise optimal (Godfray 1987; Parker and Mock 1987; Rosenheim 1993). Similarly, competition among herbivorous, gregarious insect larvae may favor a reduction in egg-clutch size of ovipositing females (Godfray and Parker 1992). Conversely, when larvae disperse readily—as is the case in many grazing insects—egg clustering or dumping may be favored (Roitberg and Mangel 1993). Under this scenario, there can be parent-offspring conflict over larval dispersal tendencies (Roitberg and Mangel 1993; Sjerps and Haccou 1994). Thus, across a wide range of invertebrate life histories, parent-offspring conflict has the potential to drive coevolution between female oviposition behavior (clutch size) and offspring behaviors that mediate group dynamics (e.g., siblicide, non-lethal aggression, and dispersal tendency).

Following Arnqvist and Rowe (2002), who described coevolutionary dynamics between traits that mediate sexual conflict, we can make three predictions regarding the evolution of traits that mediate group-size conflict. First, because antagonistic coevolution can produce rapid evolutionary change, parent-offspring conflict over group size should generate extensive interspecific variation in group-living traits. Second, because coevolutionary arms races are expected to have bouts of escalation and de-escalation (Parker 1983; Härdling 1999), trait evolution should be decoupled from phylogenetic history (minimal “phylogenetic inertia”, (Losos 1999; Arnqvist and Rowe 2002)). Third, assuming that the female optimum clutch size consistently exceeds the offspring optimum clutch size (Godfray 1987; Parker and Mock 1987; Godfray and Parker 1992; Roitberg and Mangel 1993; Godfray 1995), increases in female egg-clutch size should favor offspring behaviors that either increase competitive ability or decrease colony size (e.g., via dispersal) and vice versa, generating a negative correlation between these traits among species. Although analogous predictions have been confirmed in lineages experiencing sexual conflict (Arnqvist and Rowe 2002), they remain untested in the context of group-size conflict (but see: Mayhew 1998).

To test these predictions, we take advantage of a group of herbivorous insects for which extensive ecological data are available. The sawfly genus *Neodiprion* (order: Hymenoptera; family: Diprionidae) is a Holarctic group of approximately 50 species specialized on host plants in the family Pinaceae. Within this genus, the “*Lecontei* clade” is a monophyletic lineage of 21 eastern North American species well studied from taxonomic (Ross 1955; Ross 1961; Linnen and Smith 2012), phylogenetic (Linnen and Farrell 2007; Linnen and Farrell 2008a; Linnen and Farrell 2008b; Linnen and Farrell 2010), and life history and behavioral perspectives (Coppel and Benjamin 1965; Knerer and Atwood 1973; Knerer 1993; Costa and Louque 2001; Fowers and Costa 2003; Terbot II et al. 2017). Importantly, eastern North American *Neodiprion* have documented variation in female egg-laying behavior (Supplemental Table 1) and larval grouping behavior (Terbot II et al. 2017, Supplemental Table 1). A more detailed summary of relevant life history and ecology of *Neodiprion* sawflies can be found in the supplemental materials.

If parent-offspring conflict over group size has shaped the evolution of clutch-size traits, trait variation should match the three predictions outlined above: extensive interspecific variation, minimal phylogenetic signal, and a negative correlation between female egg-clutch size and larval grouping behavior among species. To evaluate these predictions, we combined our own observations from the field and the lab with published descriptions of female and larval behavior from 19 of the 21 eastern North American *Neodiprion* species. To gain additional insight into our data, we also examined correlations between female and larval behavior and larval colony size. If larval behaviors tend to modify initial egg-clutch size, we would expect positive correlations between larval grouping behaviors and larval colony size. Conversely, if there is minimal post-hatching modification of the initial clutch size, we would expect a stronger correlation between female clutch size and larval colony size.

## Materials and Methods

### Characterizing variation in group-size traits

We compiled data on female egg-clutch size, larval grouping behavior (aggregative tendency), and larval colony size for 19 of 21 eastern North American *Neodiprion* species (Linnen and Farrell 2008a, b). The only two species for which we were unable to obtain data were *N. cubensis* and *N. insularis*, both endemic to Cuba. Here, we briefly describe how we compiled data for each trait. Additional details are provided in the Supplemental Tables 1-4.

To estimate egg-clutch size for each species, we combined field observations, laboratory assays, and published descriptions. Field observations took place in 2002, 2015, and 2016 and consisted of counts of egg scars on needles found in close proximity to early instar larvae that were collected and reared to later instars or adults for identification. Clutch sizes were estimated from the number of egg scars found on individual branch termini. To supplement our field observations, we used laboratory assays to obtain clutch-size estimates for several species. In each assay, we released a lab-reared, virgin female into a mesh cage (35.6 x 35.6cm x 61cm) containing four *Pinus* seedlings. To obtain an average clutch size for each female, we recorded the number of eggs laid in each discrete clutch (i.e., a single branch terminus on a single seedling). For these assays, females were reared via methods described elsewhere (Harper et al. 2016) and provided seedlings from the *Pinus* species on which they (or their wild-caught progenitors) were originally collected. Finally, to supplement our own clutch-size data, we surveyed the literature and included published data from which we could infer mean clutch size. In total, we obtained clutch-size data for 16 of the 19 *Neodiprion* species included in our study (Supplemental Tables 1-2).

To measure larval aggregative tendency, we used a behavioral assay described in Terbot II et al. 2017. Briefly, we spaced five larvae of a particular species equidistantly along the perimeter of a 14.5 cm diameter petri dish and recorded their behavior for 90 minutes using a Lenovo Ideapad laptop and either a Logitech or Microsoft webcam. We then used the program Video Image Master Pro (A4Video 2016) to extract individual video frames and a custom Java application to extract one image still per every 3 minutes of video (Terbot II et al. 2017). For each extracted image, we measured all pairwise distances between each of the five larval heads. To provide an acclimation period for the larvae to adjust to the petri dish arena, we excluded the first 12 images (36 minutes) of each video (Terbot II et al. 2017) using Microsoft Excel 2013 (Microsoft 2012) and a custom Perl script. This processing pipeline produced a single mean pairwise larval distance measure for each video. Larger mean pairwise distances correspond to a greater average distance between larvae and therefore a lower “larval aggregative tendency” (i.e., a greater tendency to leave the group); smaller values correspond to a smaller average distance and a higher “larval aggregative tendency”. All larvae used in these assays were collected in the field, then reared in the lab on their natal host species via standard rearing protocols until they reached a suitable size for videotaping. To reduce developmental noise in aggregative behavior, only late-instar, feeding larvae were assayed (Terbot II et al. 2017). Collection information and other details regarding the aggregative tendency assays are included in Supplemental Tables 3-4. In total, we measured larval aggregative tendency in 19 eastern North American *Neodiprion* species.

To obtain estimates of larval colony size for each species, we combined our own field observations with published descriptions of *Neodiprion* colony size. Field observations, which were recorded between 2002-2004 and 2013-2016, consisted of approximate counts of the number of larvae found in each collected colony. Colonies were located in the field via a combination of visual searching and the use of beating sheets (i.e., tapping on branches and catching dislodged larvae on a large, square piece of fabric). With this combination of methods, we were able to collect both large, conspicuous colonies and isolated individuals. We considered sawflies to be in the same colony if they formed a distinct cluster of larvae (e.g., on the same branch tip) that was spatially isolated from other such groups. We also noted when larvae were collected as isolated individuals (i.e., colony size = 1). To supplement these field data, we included observations from published papers. In total, we obtained group-size data for 19 *Neodiprion* species.

To ensure that our data were normally distributed prior to statistical analysis, we applied a van der Waerden rank-based inverse normal transformation to all three datasets. We also multiplied each value in the aggregative tendency dataset by -1 to ensure that for all three datasets, higher values always corresponded to “large group size” phenotypes (larger egg-clutches, higher aggregative tendency, larger larval colony sizes). We used Microsoft Excel 2013 for all transformations. To determine whether species differed in trait values after accounting for variation within species, we used JMP 12 (SAS Institute 2016) to perform an analysis of variance (ANOVA) on each of the three datasets. Finally, for use in regression analyses, we calculated mean values for each species and trait. The final data set used in the analysis can be found in [REMOVED FOR DOUBLE BLIND REVIEW].

### Evaluating phylogenetic signal for group-size traits

To evaluate the extent to which each group-size trait exhibits phylogenetic signal, we estimated Pagel’s λ for each trait using BayesTraits v 3.0 (Meade and Pagel 2017). Pagel’s λ is a scaling parameter that describes how well phylogenetic relationships predict trait covariance among species, with λ=1 corresponding to a model in which a trait evolves via Brownian motion along a phylogeny (i.e., strong phylogenetic signal) and λ=0 corresponding to a star-phylogeny model in which a trait evolves independently for each species (Pagel 1999). For each trait, we used BayesTraits to estimate the likelihood of our data under the “Continuous: Random Walk (Model A)” and three different λ values (λ=0, λ=1, estimated λ). Then, to evaluate the significance of the observed phylogenetic signal, we performed likelihood ratio tests (LRTs; LR = -2ΔlnL; LR is χ^2^-distributed with 1 d.f.) in which we compared the likelihood of models in which Pagel’s λ was fixed (λ=1 or λ=0) to more complex models in which λ was estimated from the data. To account for phylogenetic uncertainty, we performed LRTs for each tree in sample of 15,000 trees from the posterior distribution of a Bayesian analysis described in Linnen and Farrell 2008a. Prior to analysis, branch lengths on each tree were scaled to time using r8s version 1.71 [58], as described in Linnen and Farrell 2010. To account for variation in sample size among traits (N = 16 for egg-clutch size, N = 19 for aggregative tendency and larval colony size), we repeated this analysis on reduced (N = 16) aggregative-tendency and larval-group-size datasets.

### Evaluating relationships between group-size traits

Because evidence for phylogenetic signal was mixed (see results), we evaluated correlations between group-size traits both with and without phylogenetic correction. For uncorrected analyses, we performed correlation analyses and linear regressions in JMP 12. To test the prediction that conflict generates correlated evolution between parent and offspring traits, we estimated Pearson’s correlation coefficient I and evaluated its significance via comparison to Student’s t-distribution. To determine which traits contribute to larval colony size, we used linear regression to model the relationship between larval colony size (dependent variable) and different combinations of explanatory variables (egg-clutch size, larval aggregative tendency, and their interaction). To compare these models, we calculated the Aikaike information criteria (AIC_C_) for all possible models. We then calculated the AIC_C_ difference between each model and the model with the lowest AIC_C_ score. We also calculated Aikaike weights and evidence ratios for each model and for the subset of models that had AIC_C_ differences less than or equal to 5. We used JMP 12 to calculate the AIC_C_ scores, generate model parameter values, evaluate significance of specific model parameters, and calculate model significance compared to a null model. To calculate AIC_C_ differences, Aikaike weights, and evidence ratios, we used Microsoft Excel 2013.

To account for possible phylogenetic constraints on trait evolution, we repeated correlation and regression analyses in BayesTraits v3.0. First, to evaluate the relationship between egg-clutch size and aggregative tendency, we computed the likelihood of the data under the “Continuous: Random Walk (Model A)” model with correlation either assumed or set to zero. To evaluate the significance of the correlation, we used a LRT to compare the models with and without the correlation. Second, to evaluate the contribution of egg-clutch size and aggregative tendency to larval colony size, we used the “Continuous: Regression” model (Organ et al. 2007), with colony size as the dependent variable and egg-clutch size, larval aggregative tendency, and egg-clutch size + larval aggregative tendency as the explanatory variables. To evaluate the significance of the regression models, we performed LRTs that compared models with and without the explanatory variables. For all correlation and regression analyses, we set λ=1. As described above, we accounted for phylogenetic uncertainty by performing all analyses on a set of 15,000 ultrametric trees sampled from the posterior distribution of a previous MCMC analysis (Linnen and Farrell, 2008a; Linnen and Farrell 2010).

## Results

### Variation in group-size traits

Egg-clutch size, larval aggregative tendency, and larval colony size varied both between and within species (Fig. 1). Although there was substantial intraspecific variation in some traits/species, ANOVAs revealed significant interspecific variation in all three traits (egg-clutch size: *F*_15, 59_ = 4.8850, p < 0.0001; larval aggregative tendency: *F*_18, 377_ = 5.3260, p < 0.0001; larval colony size: *F*_18, 360_ = 17.5712, p < 0.0001;)

**Figure 1.**
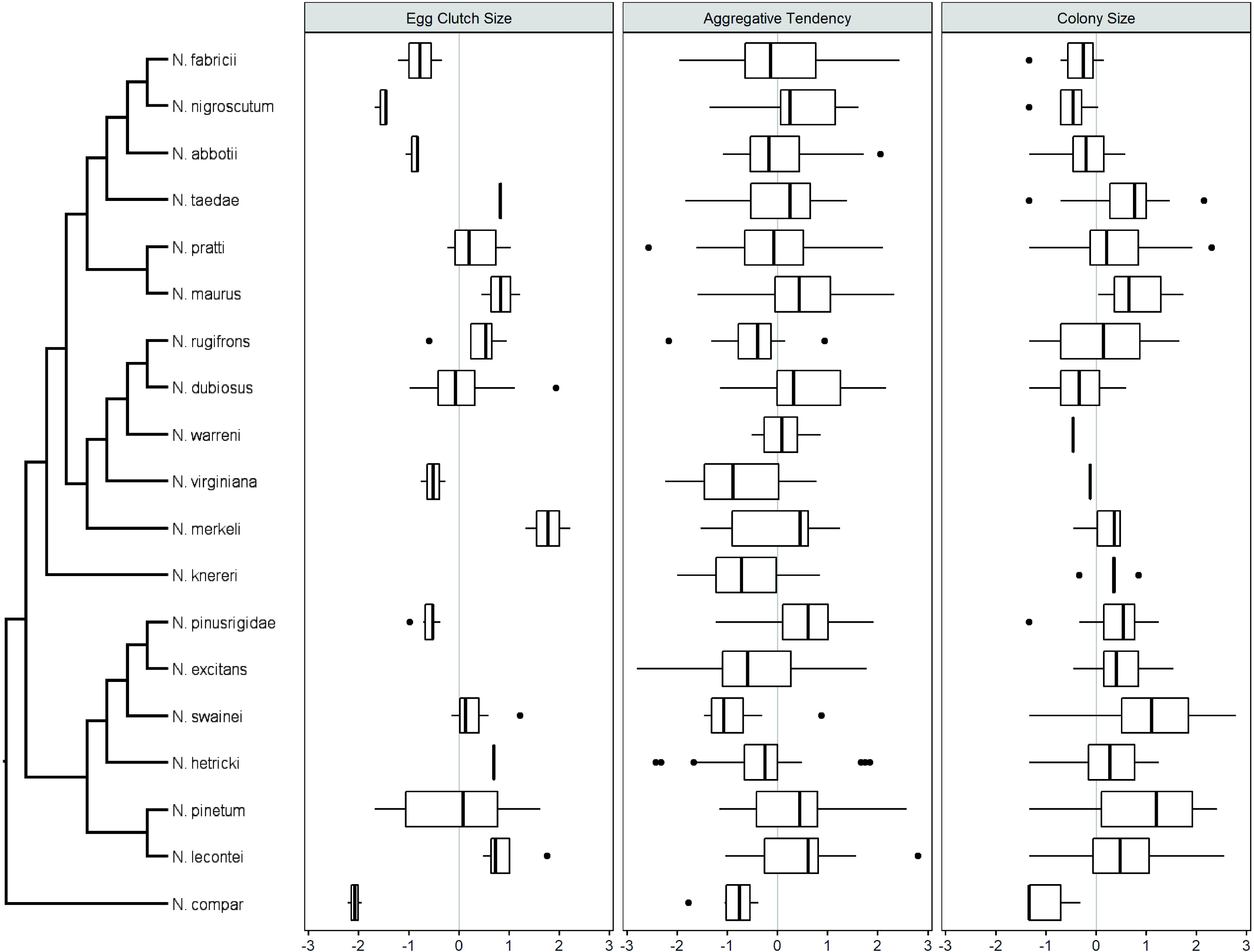
*Neodiprion* species vary in multiple group-size traits. Phylogenetic relationships for 19 eastern North American *Neodiprion* are from (Linnen and Farrell 2008a; Linnen and Farrell 2010). Box plots depict intraspecific variation for egg-clutch size, larval aggregative tendency, and larval colony size. For all three traits, there is significant interspecific variation (ANOVAs, P<0.001).

### Phylogenetic signal in group-size traits

Overall, we found strong evidence of phylogenetic signal for larval colony size, weak evidence for egg-clutch size, and no evidence for larval aggregative tendency. For larval colony size, the mean estimated λ across 15,000 trees was close to 1 (λ=0.96 ± 0.05 s.d.; Fig. 2). Moreover, while 58% of the trees rejected the model in which λ=0 (at α=0.05), only 0.03% of the trees rejected the λ=1 model. Although λ estimates for egg-clutch size were also high (mean estimated λ=0.83 ± 0.37 s.d.; Fig. 2), none of the trees rejected any of the fixed-λ models (0% for both λ=1 and λ=0). This difference may be attributable, in part, to the smaller sample size for egg-clutch size (N = 16). In support of this interpretation, we also failed to reject fixed-λ models when we repeated the larval colony size analysis using only those taxa present in the female clutch size analysis (estimated λ=0.84 ± 0.10 s.d., 0% reject λ=1, 0.17% reject λ=0). In contrast, aggregative tendency consistently rejected models in which λ=1 (100%), but not λ=0 (0%), regardless of sample size. For this trait, the mean estimated λ was 0 (± 0.00 s.d.; Fig. 2).

**Figure 2.**
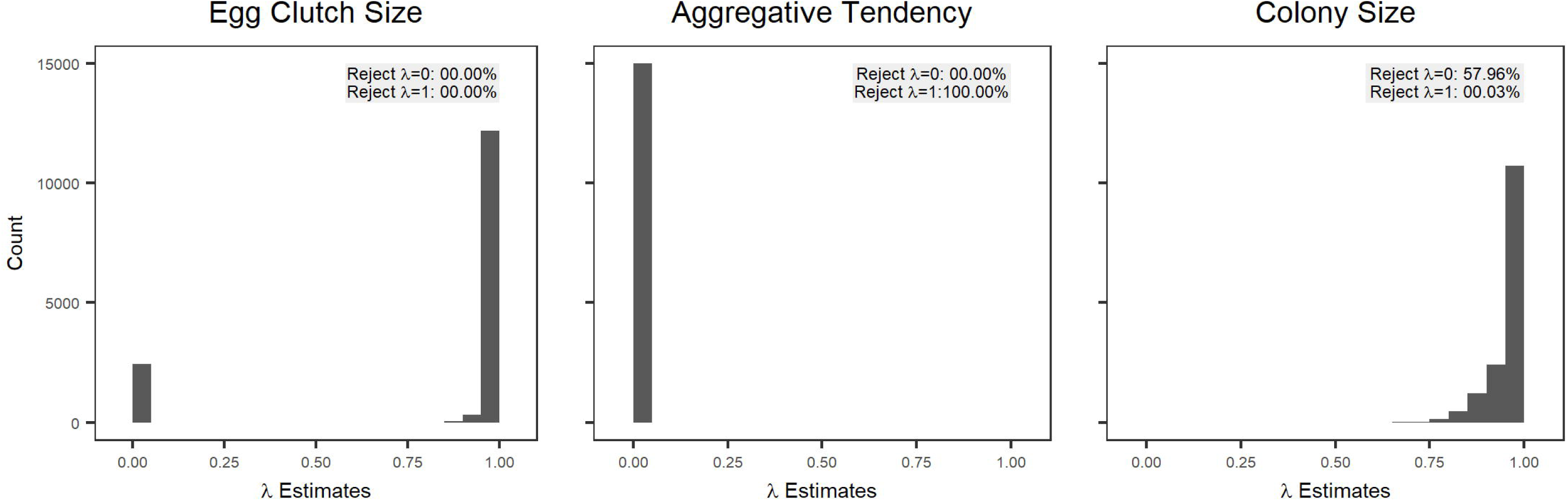
Egg-clutch size has a weak phylogenetic signal; aggregative tendency has no evidence of a phylogenetic signal; and larval colony size has a strong phylogenetic signal. Histograms of estimated lambda values across 15,000 trees sampled from the posterior distribution of a Bayesian analysis (Linnen and Farrell 2008a; Linnen and Farrell 2010) for the three traits of interest: egg-clutch size, aggregative tendency, and colony size. Noted on each histogram is the percentage of trees that rejected the fixed lambda models of λ=0 and λ=1.

### Relationships between group-size traits

Regardless of whether or not we corrected for phylogeny, we found no evidence of a correlation between egg-clutch size and aggregative tendency (non-phylogenetic: r = 0.0255, p = 0.5547, Fig. 3A; phylogenetic: estimated r = -0.0733 ± 0.0495 s.d., 0% reject null, Fig. 3D). Likewise, we found no evidence that larval aggregative tendency predicted larval colony size across species (non-phylogenetic: *F*_1,17_ = 0.0216, *R*^2^= 0.0013, p = 0.8850, Fig. 3C; phylogenetic: estimated *R*^2^ =0.0870 ± 0.0289, 0.005% reject null, Fig. 3F). In contrast, our regression analyses consistently indicated a relationship between egg-clutch size and larval colony size (non-phylogenetic: *F*_1,14_ = 13.8751, *R*^2^ = 0.4978, p = 0.0023, Fig. 3B; phylogenetic: estimated *R*^2^ =0.4997 ± 0.0356, 100% reject null, Fig. 3E). Importantly, these relationships were in the predicted direction: larger egg-clutches consistently produced larger larval groups.

**Figure 3.**
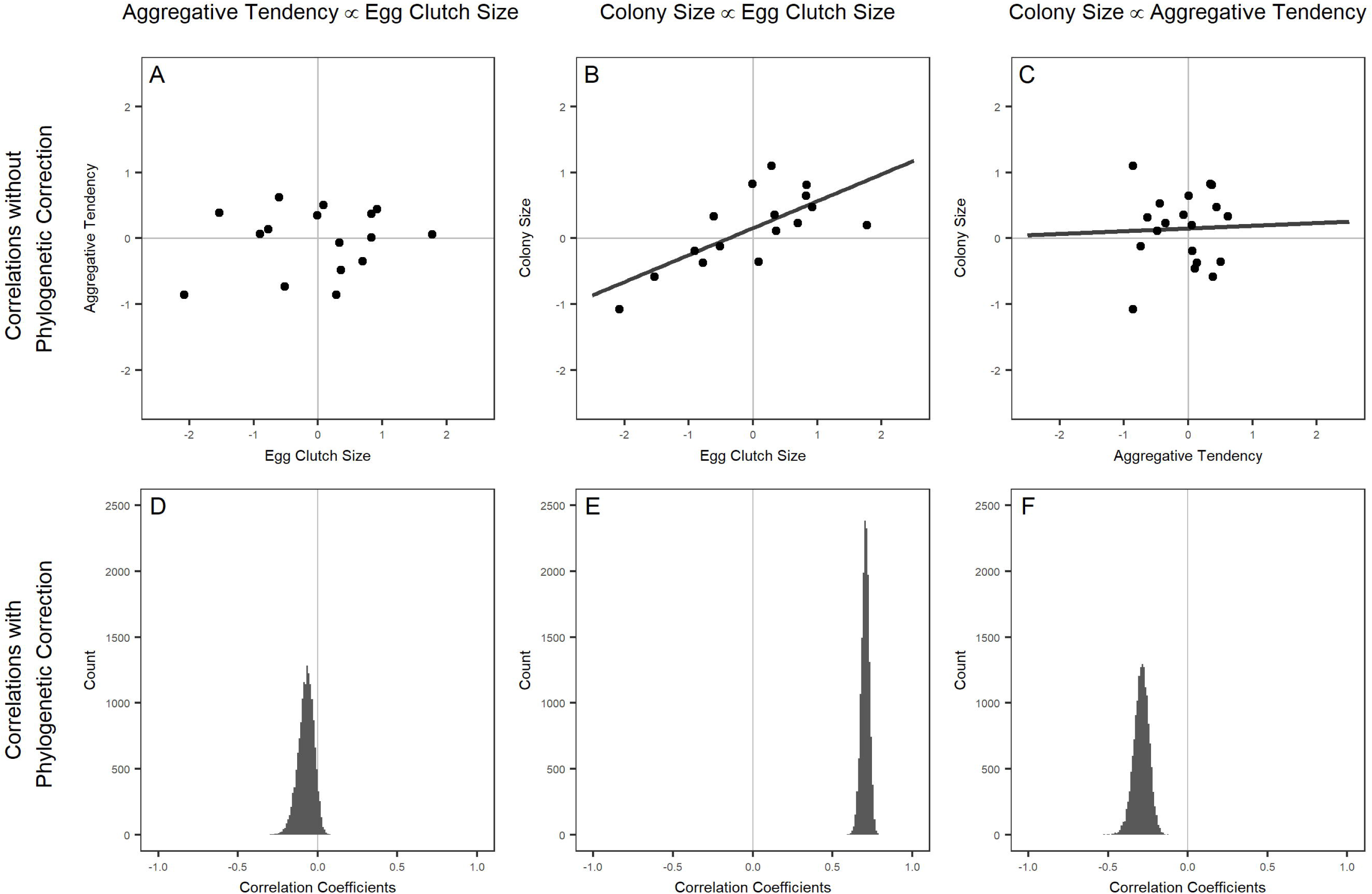
Larval aggregative tendency is not correlated with egg-clutch size and does not influence larval colony size; however, egg-clutch size predicts larval colony size. Panels A-C represent analyses lacking phylogenetic correction: egg-clutch size vs. larval aggregative tendency (correlation test, A), egg-clutch size vs. larval colony size (regression analysis, B), and aggregative tendency vs. larval colony size (regression analysis, C). Panels D-F are histograms of phylogenetically corrected correlation coefficients between traits estimated for 15,000 trees from the posterior distribution of a Bayesian analysis (Linnen and Farrell 2008a; Linnen and Farrell 2010): egg-clutch size vs. larval aggregative tendency (correlation test, D), egg-clutch size vs. larval colony size (regression analysis, E), and aggregative tendency vs. larval colony size (regression analysis, F).

Of the eight possible statistical models incorporating some combination of the explanatory traits, egg-clutch size and aggregative tendency, and their interaction; the best fitting model with the lowest AIC_C_ score was egg-clutch size alone (AIC_C_ = 23.77864; p = 0.0023 compared to a null model). The next most likely model used egg-clutch size and the interaction between egg-clutch size and aggregative tendency as explanatory variables (AIC_C_ = 25.92652, p = 0.0062 compared to a null model). The third most likely model used egg-clutch size and aggregative tendency as explanatory variables (AIC_C_ = 27.41347, p = 0.0114 compared to a null model). All other statistical models had Δ AIC_C_ scores greater than 5. The p-values for comparisons to a null model, AIC_C_ scores, Δ AIC_C_ scores, Aikaike weights and evidence ratios for all statistical models are available in Table 1.

**Table 1.** Egg-clutch size is a better predictor of larval colony size than larval aggregative tendency. Results for non-phylogenetically corrected model comparisons. In the “Model” column, “C” refers to “colony size”, “E” to “egg-clutch size”, and “A” to “aggregative tendency”. The columns contain the p-values for each model compared to the null model (colony size alone), AICc are the Akaike information critera corrected for sample size, ΔAICc are the differences between AIC scores from the best model (C ∼ E), w_*i*_ are the Akaike weights for each model, and ER_*i*_ are the evidence ratios of each model compared to the best model (C ∼ E).

| Model | p-value | AICc | $\Delta AICc$ | $w_i$ | $ER_i$ |
| --- | --- | --- | --- | --- | --- |
| $C \sim E + A + E*A$ | 0.0196 | 30.097 | 6.318 | 0.027 | 23.552 |
| $C \sim E + A$ | 0.0114 | 27.413 | 3.635 | 0.103 | 6.156 |
| $C \sim E + E*A$ | 0.0062 | 25.927 | 2.148 | 0.216 | 2.927 |
| $C \sim A + E*A$ | 0.4044 | 36.205 | 12.427 | 0.001 | 499.326 |
| $C \sim E*A$ | 0.1707 | 32.575 | 8.797 | 0.008 | 81.315 |
| $C \sim E$ | 0.0023 | 23.779 | 0 | 0.631 | 1 |
| $C \sim A$ | 0.6593 | 34.567 | 10.789 | 0.003 | 220.151 |
| C | . | 31.721 | 7.942 | 0.012 | 53.036 |

| Model | p-value | AICc | $\Delta AICc$ | $w_i$ | $ER_i$ |
| --- | --- | --- | --- | --- | --- |
| $C \sim E + A + E * A$ | 0.0196 | 30.097 | 6.318 | 0.027 | 23.552 |
| $C \sim E + A$ | 0.0114 | 27.413 | 3.635 | 0.103 | 6.156 |
| $C \sim E + E * A$ | 0.0062 | 25.927 | 2.148 | 0.216 | 2.927 |
| $C \sim A + E * A$ | 0.4044 | 36.205 | 12.427 | 0.001 | 499.326 |
| $C \sim E * A$ | 0.1707 | 32.575 | 8.797 | 0.008 | 81.315 |
| $C \sim E$ | 0.0023 | 23.779 | 0 | 0.631 | 1 |
| $C \sim A$ | 0.6593 | 34.567 | 10.789 | 0.003 | 220.151 |
| C | . | 31.721 | 7.942 | 0.012 | 53.036 |

| Model | p-value | AICc | ΔAICc | $w_i$ | $ER_i$ |
| --- | --- | --- | --- | --- | --- |
| $C \sim E + A + E * A$ | 0.0196 | 30.097 | 6.318 | 0.027 | 23.552 |
| $C \sim E + A$ | 0.0114 | 27.413 | 3.635 | 0.103 | 6.156 |
| $C \sim E + E * A$ | 0.0062 | 25.927 | 2.148 | 0.216 | 2.927 |
| $C \sim A + E * A$ | 0.4044 | 36.205 | 12.427 | 0.001 | 499.326 |
| $C \sim E * A$ | 0.1707 | 32.575 | 8.797 | 0.008 | 81.315 |
| $C \sim E$ | 0.0023 | 23.779 | 0 | 0.631 | 1 |
| $C \sim A$ | 0.6593 | 34.567 | 10.789 | 0.003 | 220.151 |
| $C$ | . | 31.721 | 7.942 | 0.012 | 53.036 |

## Discussion

Comparative data across diverse systems suggest that antagonistic coevolution stemming from evolutionary conflicts can fuel rapid diversification of traits and species (Crespi and Nosil 2013; Carmona et al. 2015; Queller and Strassmann 2018). Such dynamics have been confirmed in the context of parent-offspring conflict in organisms providing substantial parental care (Briskie et al. 1994; Lloyd and Martin 2003). Although conflicts are also expected even in the absence of parental care (Godfray and Parker 1987; Parker and Mock 1987; Godfray and Parkre 1992; Roitbeerg and Mangel 1993; Rosenheim 1993; Sjerps and Haccou 1994), comparative studies examining such systems are lacking (but see Mayhew 1998). In this study, we have taken advantage of a genus of herbivorous insects with well-documented interspecific variation in traits that mediate group size to ask whether evolutionary dynamics are consistent with parent-offspring conflict. While we find extensive interspecific variation in all three traits and some decoupling from phylogeny, there is no evidence of coevolution between adult female oviposition behavior and larval grouping behavior. Furthermore, while larvae have the physical capacity to disperse (Benjamin 1955; Smirnoff 1960; Codella and Raffa 1995), the close correspondence between egg-clutch size and larval colony size suggests that variable aggregative behavior in the larvae does not substantially alter the initial clutch size. A discussion of possible limitations of our phenotypic datasets can be found in the supplemental materials. Here, we consider possible biological explanations for the apparent lack of strong parent-offspring conflict over group size and propose alternative explanations for interspecific variation in group-living traits.

One potential explanation for the apparent lack of correlated evolution between female clutch-size and larval grouping behavior is that populations often lack heritable variation in grouping traits that mediate conflicts over group size. However, this explanation seems unlikely because we have detected considerable interspecific variation in both oviposition traits and larval aggregative tendency, even when measured under uniform laboratory conditions (Fig. 1). Although non-genetic sources (e.g., plasticity) may explain some of this variation, experimental crosses have confirmed that some of this behavioral variation is attributable to genetic differences between populations and species (Knerer and Atwood 1972; Bendall et al 2017). That said, even with genetic variation for adult and larval behaviors, physiological limits on some traits could constrain trait evolution in response to parent-offspring conflict over group size. For example, sawfly females typically emerge from cocoons with their full complements of eggs (Coppel and Benjamin 1965), placing an upper limit on maximum clutch size. This upper limit could prevent egg-clutch sizes from becoming large enough to generate strong selection on dispersal tendencies, thereby limiting the potential for repeated bouts of reciprocal evolutionary change.

Assuming that populations could respond to selection stemming from parent-offspring conflict, another explanation for a lack of correlated evolution is that conflict is too weak or idiosyncratic to produce such a pattern. For example, many theoretical studies on group-size conflict assume that larval fitness is optimized at a group size of n = 1 (Parker and Mock 1987; Godfray and Parker 1992; Roitberg and Mangel 1993; but see Godfray and Ives 1988), ignoring possible advantages to larval grouping (e.g., an Allee effect (Courchamp et al. 1999)). By contrast, potential benefits of group-living are well documented (Seymour 1974; Bertram 1978; Young and Moffett 1979; Tsubaki and Shiotsu 1982; Pulliam and Caraco 1984; Joos et al. 1988; Stamp and Bowers 1990; Codella and Raffa 1993; Klok and Chown 1999; Despland and Le Huu 2007; Fletcher 2009; McClure et al. 2011a; McClure et al. 2011b; McClure et al. 2013; Campbell and Stastny 2015). In *Neodiprion*, it has been hypothesized that group-living helps larvae overcome host defenses (Ghent 1960, but see Kalin and Knerer 1977). If benefits of group-living consistently outweigh costs in *Neodiprion*, parent-offspring conflict over group size could be minimal.

Overall, our data suggest that parent-offspring conflict is not the predominant force shaping interspecific variation in female clutch size and larval grouping behavior. Another potential explanation for among-species variation in these traits is that it accumulated via genetic drift as species diverged from common ancestors. Under this scenario, however, we would expect a strong phylogenetic signal for grouping traits. By contrast, our data indicate that traits are at least partially (female clutch-size) or completely (larval aggregative tendency) decoupled from phylogeny. Therefore, we now consider other possible selection pressures that may shape intra- and interspecific variation in these traits. Ultimately, testing these hypotheses will require a combination of experiments that verify targets and agents of selection and comparative analyses that evaluate the contribution of different selective mechanisms to genus-wide variation.

For an adult female, optimal clutch size depends on the distribution and quality of available host plants, the physiological condition of the female, and the relationship between clutch size and offspring survival (Skinner 1985; Mangel 1987; Tostowaryk 1972). Variation in any of these factors could favor different oviposition strategies in different species. For example, *Neodiprion* species vary in their preferred oviposition hosts (Knerer and Atwood 1972; Knerer and Atwood 1973; Linnen and Farrell 2010; Bendall et al. 2017). If preferred hosts differ in their patchiness, different oviposition strategies may be favored. Interspecific variation in clutch size may also arise if the relationship between clutch size and larval survival varies among species. For example, some *Neodiprion* species have aposematic coloration (Tostwaryk 1972; Linnen et al. 2018). When aposematic signals are used, group-living can enhance the efficacy of those signals (Gamberale and Tullberg 1998; Hatle and Salazar 2001; Riipi et al. 2001). Anecdotally, *Neodiprion* with brighter coloration (e.g., *Neodiprion lecontei*) tend to have larger colony sizes than more cryptically colored species (e.g., *N. compar*). However, additional comparative work is needed to determine whether group-living correlates with conspicuousness in *Neodiprion*.

For larvae, although aggregative tendency does not appear to impact larval group size, it may nevertheless play an important role in larval survival. By controlling the density of larvae within groups, variation in aggregative tendency can impact disease transmission rates, microclimate, and host physical and chemical defenses. First, because disease transmission occurs primarily through physical contact, reduced disease transmission is one possible advantage of having a diffuse larval group (i.e., low aggregative tendency). Notably, nucleopolyhedrovirus and other infectious diseases can be a major source of mortality for sawfly larvae and gregarious larvae of ecologically similar species (Bird 1955; Mohamed et al 1985; Young and Yearian 1987; Hochbereg 1991; Fletcher 2009). Second, through managing the larval group’s density, larval aggregative tendency could also be a mechanism through which group-living species optimize their micro-environment (Seymour 1974; Joos et al. 1988; Codella and Raffa 1993; Klok and Chown 1999; Fletcher 2009; McClure et al. 2011a). For example, compared to warm and wet environments, cold or dry environments may favor tighter clustering of larval groups. Third, variation in pine needle morphology and defensive chemistry among pine species may favor different larval densities. When feeding in groups, sawfly larvae form a characteristic “ring” around a needle near the tip and, as a group, consume the needle tissue down to the fascicle (Ghent 1960). However, the ability to form a stable feeding ring may decrease on thin- and long-needled hosts. Reduced aggregative tendency may prevent larvae from overloading thin needles unable to bear their weight. By contrast, thick and resinous needles may favor increased aggregative tendency to ensure a sufficient number of larvae to establish feeding sites and distribute host resins (Ghent 1960).

## Conclusions

Comparative analysis is a powerful tool for testing and honing intuitions derived from evolutionary theory. For example, although theoretical work suggests that small, gregarious broods of parasitoids are evolutionary unstable and transitions from solitary to gregarious states should be rare (Godfray 1987), clutch-size data from parasitoid wasps suggests that such transitions have occurred many times independently (Mayhew 1998). Analogously, although theory demonstrates the potential for parent-offspring conflict over group size in plant-feeding insects (Roitberg and Mangel 1993), our data suggest that this conflict is not sufficiently strong or consistent to drive correlated evolution of adult and larval grouping traits in pine sawflies. To be clear, these findings do not rule out parent-offspring conflict entirely. Rather, they suggest that when parental care is minimal, parent-offspring conflict is not a major driver of long-term patterns of phenotypic change. Although these trends are intriguing, more comparative data are needed to evaluate whether some forms of conflict are more likely than others to leave a detectable phenotypic footprint in phylogenetic comparative analyses.

## Supporting information

Supplemental discussions of life history and study caveats as well as details on data and specimen collection.

