## Supplemental discussions of life history and study caveats as well as details on data and specimen collection. for "Mother knows best: maternal egg-clutch size predicts larval group size in pine sawflies (genus *Neodiprion*)"

**Supplementary Materials**

Neodiprion *sawflies*: *life history description and utility as a model for parent-offspring conflict*

To provide context for our comparative analysis of *Neodiprion* clutch-size traits, we provide relevant life history details (reviewed in Coppel and Benjamin 1965; Knerer and Atwood 1973; Wilson et al. 1992; Knerer 1993). Adult females emerge from cocoons with a full complement of mature eggs, find a suitable host, and attract males via a powerful pheromone. Shortly after mating, females use their saw-like ovipositors to embed their eggs within pine needles. While females of some species tend to lay their full complement of eggs on a single branch terminus, females of other species seek out multiple branches or trees for oviposition. Overall, female oviposition behavior is highly species-specific and, in some cases, diagnostic (Ghent 1959). Adult *Neodiprion* are non-feeding and short-lived (~2-4 days), dying soon after mating and oviposition. After hatching from eggs, *Neodiprion* larvae of many species form feeding aggregations that remain intact to varying degrees across 4-7 feeding instars, depending on the sex and the species. As larvae defoliate pine branches, they migrate to new branches and sometimes to new host trees (Benjamin 1955, Smirnoff 1960). During these migrations, colonies may undergo fission and fusion events (Codella and Raffa 1993, Codella and Raffa 1995, Costa and Loque 2001). Thus, while initial colony size corresponds closely to egg-clutch size, larvae are highly mobile and their dispersal behavior has the potential to substantially alter colony size (Codella and Raffa 1995).

Beyond having a variable and well-documented natural history, *Neodiprion* provides an excellent test case for examining coevolution of female egg-laying and larval grouping behaviors because, as is likely the case for many insects, feeding in groups could confer both costs and benefits to the larvae (Codella and Raffa 1995, Heitland and Pschorn-Walcher 1993). On the one hand, larvae living in large groups compete for access to pine needles (Prokopy *et al* 1984). Also, because large colonies quickly defoliate branches, larvae must expend energy and risk exposure to predators to reach new branches or trees (Benjamin 1955, Smirnoff 1960, Teras 1982, Costa & Louque 2001, Flowers and Costa 2003). Compared to small colonies, large colonies may be more conspicuous to predators and parasitoids (Lyons 1962, Olofsson 1986) and more susceptible to viral infection (Mohamed et al. 1985; Young and Yearian 1990). On the other hand, grouping is hypothesized to confer numerous benefits to larval colonies. For example, in early instars, grouping is thought to be essential to establishing feeding sites on tough pine needles (Ghent 1960, but see Kalin and Knerer 1977). Also, living in large groups may decrease per capita predation risk (Hamilton 1971; Codella and Raffa 1995) or enhance aposematic signalling (Tostowaryk 1972, Sillen-Tullberg & Leimar 1988, Linnen et al. 2018).

*Limitations of our phenotypic data*

Although the main findings of our study are robust to analysis method and phylogenetic uncertainty, there are limitations of the three phenotypic datasets that should be revisited with additional laboratory assays and field studies. First, our egg-clutch size dataset had more missing data than the other datasets, which reduced our power to detect phylogenetic signal and correlated evolution with other traits. Nevertheless, we did detect a strong relationship between egg-clutch size and larval grouping behavior (Fig. 3B, Table 1). The egg-clutch data also relied more heavily on literature records than other datasets (Sup. Table 2). By and large, these literature records contain detailed descriptions of oviposition behavior and conform well to our own observations. Also, where possible, we supplemented published egg-clutch size data with our own field observations and lab studies. For field observations, we assumed that eggs found on a single branch terminus were laid by a single female. In support of this assumption, ovipositing females of some species appear to produce an oviposition deterrent that may prevent multiple-female clutches (Tisdale & Wagner 1991). That said, "contagious oviposition" at the level of host trees has been reported for some *Neodiprion* species (Wilson 1975; Codella and Raffa 1995) and there are anecdotal reports of multiple females ovipositing in a single branch terminus as well (Warren & Coyne 1958; Codella and Raffa 1995). Thus, it is possible that some of our field egg-clutch observations over-estimated female clutch size. Our laboratory measurements may have also over-estimated clutch size due to a small test arena and limited availability of branch termini.

Second, in contrast to the egg-clutch data, all aggregative tendency data were obtained from laboratory assays. While the use of a uniform quantitative assay likely reduced noise in our behavioral data, there are several limitations of this assay (see Terbot *et al* 2017). Most relevant for this study, the small, host-free test arena used in this assay may have simultaneously increased larval tendency to disperse (e.g., in search of host foliage) and reduced our power to detect differences in larval aggregative tendency (e.g., due to the inability to leave the small arena). Despite these limitations, a previous study demonstrated that this assay is sufficient to detect behavioral differences between different *Neodiprion* species and between different developmental stages (feeding versus wandering larvae) of a single species (Terbot *et al* 2017). Beyond assay design, another limitation of our data is that we did not control for sex, which is difficult to determine from larval morphology (Wilkinson 1971). Although it is currently unknown whether sexes differ in their larval aggregative tendency, sex-based differences–which have been documented for other larval behaviors in pine sawflies (Lindstedt et al. 2018) – could potentially introduce noise into our data. Finally, there are additional larval behaviors that we have not measured that could also impact colony size, such as trail following during migration between feeding sites (Costa & Louque 2001, Flowers & Costa 2003).

Third, the majority of our colony size data points came from direct field observations obtained from all developmental stages (different feeding larval instars). Because developmental variation in gregarious behaviors is well documented in pine sawflies and other insects (Damman 1991, McClure & Despland 2011, Klok & Chown 1999, Furniss & Dowden 1941, Hetrick 1956, Terbot et al. 2017), our inclusion of different instars likely introduced noise into our larval colony size dataset. Unfortunately, we did not have sufficient field data to compare larval colony size results from different developmental stages. Nevertheless, the strong correlation we observed between larval colony size and egg-clutch size (Fig. 3B, Table 1) suggests that group size is relatively stable across development. Another potential issue with our field observations stems from detection bias. Because larger groups of larvae are easier to see, they may be overrepresented in our data and bias our colony-size estimates. However, this potential source of bias is mitigated to some extent by our use of beating sheets to locate smaller colonies and solitary individuals, both of which were well represented in our dataset. Moving forward, the relationship between egg-clutch size, larval grouping behaviors, and larval group size could be further clarified by carefully tracking individual egg-clutches from hatching to cocooning (e.g., as with *N. lecontei* in Costa & Louque 2001 and Flowers & Costa 2003) in many different *Neodiprion* species.

Although our analysis of grouping behavior variation would clearly benefit from additional phenotypic measurements, the existing dataset produces robust results that are unlikely to be explained by biases in our data collection. Specifically, despite considerable interspecific variation in larval aggregative behaviors, our data suggests that this variation is decoupled from both maternal oviposition behavior and larval group size.

Supplemental Table 1. Table of literature sources for colony size and egg clutch size data.

| Citation | Species | Colony Size Records | Egg Clutch Size Records |
|---|---|---|---|
| Atwood, 1961 | *N. compar* | 1 | |
| Atwood & Peck, 1943 | *N. abbotii* | | 1 |
| Atwood & Peck, 1943 | *N. compar* | | 1 |
| Baker, 1972 | *N. lecontei* | | 1 |
| Baker, 1972 | *N. taedae* | | 1 |
| Becker, 1965 | *N. rugifrons* | | 1 |
| Becker, 1965 | *N. dubiosus* | 2 | 2 |
| Becker, 1965 | *N. nigroscutum* | 1 | |
| Becker & Benjamin, 1964 | *N. swainei* | | 1 |
| Becker & Benjamin, 1967 | *N. nigroscutum* | 2 | 1 |
| Codella & Raffa, 2002 | *N. lecontei* | | 1 |
| Ghent, 1955 | *N. pratti* | | 1 |
| Ghent & Wallace, 1958 | *N. swainei* | | 1 |
| Hetrick, 1941 | *N. taedae* | | 1 |
| Hetrick, 1956 | *N. abbotii* | | 1 |
| Hetrick, 1956 | *N. hetricki* | | 1 |
| Jansons, 1966 | *N. compar* | 1 | |
| Knerer, 1984 | *N. pratti* | | 1 |
| Knerer, 1990 | *N. maurus* | | 1 |
| Lyons, 1964 | *N. swainei* | | 4 |
| Middleton, 1921 | *N. lecontei* | | 1 |
| Rauf & Benjamin, 1980 | *N. maurus* | | 1 |
| Rauf & Benjamin, 1980 | *N. pinetum* | | 1 |
| Schedl, 1938 | *N. dubiosus* | | 1 |
| Smirnoff, 1960 | *N. swainei* | 2 | |
| Tostowaryk, 1972 | *N. pratti* | | 1 |
| Tostowaryk, 1972 | *N. swainei* | | 1 |
| Wilkinson, 1961 | *N. rugifrons* | 3 | 1 |
| Wilkinson, 1965 | *N. merkeli* | | 1 |
| Wilkinson, 1971 | *N. merkeli* | | 1 |
| Wilkinson, 1978 | *N. pratti* | | 1 |
| Wilkinson, Becker, & Benjamin, 1966 | *N. rugifrons* | 1 | 1 |
| Wilson, 1971 | *N. pratti* | 1 | 1 |
| Wilson, 1977 | *N. nigroscutum* | 1 | |
| Wilson, 1977 | *N. abbotii* | | 1 |

Supplemental Table 2. Summary of sample sizes and sources of data for colony size, egg clutch size, and aggregative tendency data sets

| Species | Colony Size | | | Egg Clutch Size | | | | Aggregative Tendency |
|---|---|---|---|---|---|---|---|---|
| | Field Observations | Literature References | Total | Field Observations | Lab Behavior | Literature References | Total | Lab Behavior |
| *N. abbotii* | 9 | 0 | 9 | 0 | 0 | 3 | 3 | 19 |
| *N. compar* | 62 | 2 | 64 | 1 | 0 | 1 | 2 | 8 |
| *N. dubiosus* | 34 | 1 | 35 | 9 | 0 | 3 | 12 | 27 |
| *N. excitans* | 11 | 0 | 11 | 0 | 0 | 0 | 0 | 36 |
| *N. fabricii* | 17 | 0 | 17 | 0 | 2 | 0 | 2 | 12 |
| *N. hetricki* | 16 | 0 | 16 | 0 | 0 | 1 | 1 | 35 |
| *N. knereri* | 5 | 0 | 5 | 0 | 0 | 0 | 0 | 8 |
| *N. lecontei* | 37 | 1 | 38 | 1 | 0 | 3 | 4 | 26 |
| *N. maurus* | 19 | 0 | 19 | 0 | 0 | 2 | 2 | 22 |
| *N. merkeli* | 6 | 0 | 6 | 0 | 0 | 2 | 2 | 10 |
| *N. nigroscutum* | 14 | 4 | 18 | 1 | 0 | 2 | 3 | 8 |
| *N. pinetum* | 9 | 0 | 9 | 0 | 13 | 1 | 14 | 22 |
| *N. pinusrigidae* | 9 | 0 | 9 | 7 | 0 | 0 | 7 | 42 |
| *N. pratti* | 46 | 1 | 47 | 0 | 3 | 5 | 8 | 46 |
| *N. rugifrons* | 32 | 4 | 36 | 1 | 0 | 3 | 4 | 16 |
| *N. swainei* | 16 | 2 | 18 | 0 | 0 | 7 | 7 | 11 |
| *N. taedae* | 19 | 0 | 19 | 0 | 0 | 2 | 2 | 23 |
| *N. virginiana* | 1 | 0 | 1 | 0 | 2 | 0 | 2 | 19 |
| *N. warreni* | 2 | 0 | 2 | 0 | 0 | 0 | 0 | 6 |

Supplemental Table 3. Collection information and summary of data points per site.

| Collection identifier | Species | Number of Colony Size Observations | Number of Field Egg Clutch Observations | Number of Aggregative Tendency Videos | Number of In-Lab Oviposition Cages | Date of Collection | Latitude | Longitude |
|---|---|---|---|---|---|---|---|---|
| 001-04 | N. abbotii | 1 | | | | 04/23/04 | 30.2908 | -82.0749 |
| NS013 | N. abbotii | 1 | | | | 09/01/13 | 35.1310 | -85.4000 |
| NS014 | N. abbotii | 1 | | | | 09/01/13 | 35.5670 | -84.4630 |
| NS021 | N. abbotii | 1 | | 1 | | 03/19/14 | 29.7797 | -84.7011 |
| NS027 | N. abbotii | 1 | | | | 04/28/14 | 35.4506 | -80.5947 |
| 122-04B | N. abbotii | 1 | | | | 07/20/14 | 35.2588 | -81.3600 |
| NS057 | N. abbotii | | | 2 | | 08/03/14 | 46.6231 | -81.4591 |
| NS072 | N. abbotii | | | 2 | | 08/18/14 | 45.1316 | -91.9247 |
| NS086 | N. abbotii | | | 1* | | 09/07/14 | 38.1722 | -83.5570 |
| NS088 | N. abbotii | | | 1* | | 09/07/14 | 38.1039 | -83.5113 |
| NS114.03 | N. abbotii | 1 | | 1 | | 04/25/15 | 36.1791 | -85.9456 |
| NS117.03 | N. abbotii | | | 4 | | 04/25/15 | 35.9419 | -86.5278 |
| NS141 | N. abbotii | | | 3 | | 05/10/15 | 37.7862 | -80.3004 |
| NS162.01 | N. abbotii | 1 | | 5 | | 07/25/15 | 44.5511 | -68.3945 |
| 147-02 | N. compar | 1 | | | | 08/08/02 | 45.9264 | -77.3256 |
| 148-02 | N. compar | 1 | | | | 08/08/02 | 45.9264 | -77.3256 |
| 149-02 | N. compar | 1 | | | | 08/08/02 | 45.9264 | -77.3256 |
| 154-02 | N. compar | 1 | | | | 08/08/02 | 45.9316 | -77.3332 |
| 158-02 | N. compar | 1 | | | | 08/08/02 | 45.9316 | -77.3332 |
| 159-02 | N. compar | 1 | | | | 08/08/02 | 45.9673 | -77.3715 |
| 194-02 | N. compar | 1 | | | | 08/11/02 | 46.1334 | -80.6873 |
| 197-02 | N. compar | 2 | | | | 08/11/02 | 46.6240 | -81.4544 |
| 203-02 | N. compar | | 1 | | | 08/11/02 | 46.6240 | -81.4544 |
| 224-02 | N. compar | 1 | | | | 08/13/02 | 48.5500 | -89.7115 |
| 234-02 | N. compar | 1 | | | | 08/14/02 | 49.7247 | -94.2481 |
| 241-02 | N. compar | 1 | | | | 08/14/02 | 49.7247 | -94.2481 |
| 255-02 | N. compar | 1 | | | | 08/15/02 | 48.6885 | -93.0019 |
| 263-02 | N. compar | 1 | | | | 08/16/02 | 48.6315 | -85.3619 |
| 265-02 | N. compar | 1 | | | | 08/16/02 | 48.5423 | -85.1882 |
| 266-02 | N. compar | 1 | | | | 08/16/02 | 48.0295 | -84.6518 |
| 289-02 | N. compar | 1 | | | | 08/17/02 | 48.2203 | -82.0736 |
| 312-02 | N. compar | 1 | | | | 08/18/02 | 47.4835 | -81.8459 |
| 321-02 | N. compar | 1 | | | | 08/18/02 | 47.4849 | -81.4034 |

| | | | | | | |
|---|---|---|---|---|---|---|
| 326-02 | N. compar | 1 | | 08/18/02 | 47.6471 | -80.9150 |
| 371-02 | N. compar | 1 | | 09/25/02 | 41.8828 | -70.6385 |
| 378-02 | N. compar | 1 | | 09/25/02 | 41.8658 | -70.6576 |
| 380-02 | N. compar | 1 | | 09/25/02 | 41.8658 | -70.6576 |
| 111-04A | N. compar | 1 | | 07/19/04 | 36.1370 | -85.5031 |
| 114-04 | N. compar | 1 | | 07/19/04 | 36.0484 | -84.9640 |
| 128-04 | N. compar | 1 | | 07/22/04 | 37.1854 | -81.1285 |
| 163-04 | N. compar | 1 | | 07/29/04 | 45.4906 | -70.2360 |
| 182-04 | N. compar | 1 | | 08/14/04 | 44.5709 | -91.6352 |
| 214-04 | N. compar | 1 | | 08/17/04 | 46.5007 | -90.4621 |
| NS029 | N. compar | 1 | | 05/04/14 | 37.1015 | -77.5426 |
| 122-04A | N. compar | 1 | | 07/20/14 | 35.2588 | -81.3600 |
| NS148 | N. compar | 1 | | 06/14/15 | 33.9282 | -83.3761 |
| NS169.01 | N. compar | | 1* | 08/13/15 | 45.9315 | -77.3333 |
| NS170 | N. compar | | 1* | 08/14/15 | 46.6231 | -81.4552 |
| NS172.01 | N. compar | 1 | 1* | 08/14/15 | 47.4648 | -81.8467 |
| NS174.01 | N. compar | 1 | 2* | 08/15/15 | 48.0456 | -84.5494 |
| NS175 | N. compar | | 1* | 08/15/15 | 48.0297 | -84.6513 |
| NS176 | N. compar | 1 | 1* | 08/15/15 | 48.5437 | -85.1911 |
| NS178.01 | N. compar | 1 | 1* | 08/16/15 | 48.4161 | -89.6412 |
| NS182.01 | N. compar | 1 | 1* | 08/17/15 | 46.5089 | -90.5027 |
| NS183 | N. compar | 1 | | 08/17/15 | 46.5006 | -90.4626 |
| NS184.01 | N. compar | | 1* | 08/17/15 | 46.1149 | -90.5511 |
| NS217 | N. compar | 6 | 1 | 08/15/16 | 48.0302 | -84.6490 |
| NS218 | N. compar | 11 | 2+1* | 08/15/16 | 48.0459 | -84.5507 |
| NS220 | N. compar | 5 | 1* | 08/15/16 | 48.6312 | -85.3626 |
| NS221 | N. compar | 2 | 1* | 08/15/16 | 48.7935 | -87.0895 |
| NS217 | N. dubiosus | 2 | | 08/15/16 | 48.0302 | -84.6490 |
| 203-02 | N. dubiosus | 1 | | 08/11/02 | 46.6240 | -81.4544 |
| 210-02 | N. dubiosus | 1 | | 08/13/02 | 48.3780 | -89.5799 |
| 216-02 | N. dubiosus | 1 | | 08/13/02 | 48.3763 | -89.5844 |
| 217-02 | N. dubiosus | 1 | | 08/13/02 | 48.4156 | -89.6378 |
| 226-02 | N. dubiosus | 1 | | 08/13/02 | 48.5500 | -89.7115 |
| 245-02 | N. dubiosus | 1 | | 08/14/02 | 49.7247 | -94.2481 |
| 252-02 | N. dubiosus | 1 | | 08/15/02 | 48.6885 | -93.0019 |
| 276-02 | N. dubiosus | 1 | | 08/16/02 | 47.9763 | -84.5318 |
| 278-02 | N. dubiosus | 1 | | 08/16/02 | 47.9763 | -84.5318 |
| 280-02 | N. dubiosus | 1 | | 08/16/02 | 47.9584 | -84.2683 |
| 281-02 | N. dubiosus | 1 | | 08/17/02 | 47.9337 | -83.106 |
| 282-02 | N. dubiosus | 1 | | 08/17/02 | 48.0934 | -82.6727 |
| 284-02 | N. dubiosus | 1 | | 08/17/02 | 48.0934 | -82.6727 |
| 285-02 | N. dubiosus | 1 | | 08/17/02 | 48.0934 | -82.6727 |
| 286-02 | N. dubiosus | 1 | | 08/17/02 | 48.1851 | -82.2276 |

| | | | | | | | |
|---|---|---|---|---|---|---|---|
| 288-02 | N. dubiosus | 1 | | | 08/17/02 | 48.2203 | -82.0736 |
| 290-02 | N. dubiosus | 1 | | | 08/17/02 | 48.2203 | -82.0736 |
| 293-02 | N. dubiosus | 1 | | | 08/17/02 | 48.2203 | -82.0736 |
| 295-02 | N. dubiosus | | 1 | | 08/17/02 | 48.3052 | -81.7574 |
| 297-02 | N. dubiosus | 1 | | | 08/17/02 | 48.3651 | -81.5758 |
| 300-02 | N. dubiosus | 1 | | | 08/17/02 | 47.8084 | -81.5917 |
| 306-02 | N. dubiosus | 1 | | | 08/17/02 | 47.8084 | -81.5917 |
| 309-02 | N. dubiosus | 1 | | | 08/18/02 | 47.7059 | -81.7576 |
| 311-02 | N. dubiosus | 1 | | | 08/18/02 | 47.4835 | -81.8459 |
| 323-02 | N. dubiosus | 1 | | | 08/18/02 | 47.4849 | -81.4034 |
| 327-02 | N. dubiosus | 1 | | | 08/18/02 | 47.6471 | -80.915 |
| 329-02 | N. dubiosus | 1 | | | 08/18/02 | 47.6471 | -80.915 |
| 330-02 | N. dubiosus | 1 | | | 08/18/02 | 47.6471 | -80.915 |
| 331-02 | N. dubiosus | 1 | | | 08/18/02 | 47.6651 | -80.6184 |
| 334-02 | N. dubiosus | 1 | | | 08/18/02 | 47.6651 | -80.6184 |
| 338-02 | N. dubiosus | 1 | | | 08/19/02 | 47.4254 | -79.7427 |
| 161-04 | N. dubiosus | 1 | | | 07/29/04 | 45.4872 | -70.2462 |
| NS059 | N. dubiosus | | | 2 | 08/04/14 | 48.1717 | -80.2546 |
| NS060 | N. dubiosus | | | 4 | 08/04/14 | 47.7313 | -80.3358 |
| NS061.01 | N. dubiosus | | | 6 | 08/04/14 | 47.6537 | -81.0517 |
| NS062 | N. dubiosus | | | 4 | 08/04/14 | 47.4836 | -81.8459 |
| NS064 | N. dubiosus | | | 4 | 08/05/14 | 48.2414 | -82.3542 |
| NS066 | N. dubiosus | | | 4 | 08/05/14 | 47.8744 | -83.2854 |
| NS151.01 | N. dubiosus | | | 2 | 06/27/15 | 48.2205 | -82.0732 |
| NS151.03 | N. dubiosus | | | 1 | 06/27/15 | 48.2205 | -82.0732 |
| NS151.04 | N. dubiosus | | 1 | | 06/27/15 | 48.2205 | -82.0732 |
| NS151.05 | N. dubiosus | | 1 | | 06/27/15 | 48.2205 | -82.0732 |
| NS151.06 | N. dubiosus | | 1 | | 06/27/15 | 48.2205 | -82.0732 |
| NS151.07 | N. dubiosus | | 1 | | 06/27/15 | 48.2205 | -82.0732 |
| NS151.08 | N. dubiosus | | 1 | | 06/27/15 | 48.2205 | -82.0732 |
| NS152.01 | N. dubiosus | | 1 | | 06/27/15 | 47.4838 | -81.8460 |
| NS152.02 | N. dubiosus | | 1 | | 06/27/15 | 47.4838 | -81.8460 |
| NS154.01 | N. dubiosus | | 1 | | 06/27/15 | 46.3111 | -81.6557 |
| NS219 | N. dubiosus | 1 | | | 08/15/16 | 48.5437 | -85.1910 |
| 175-03 | N. excitans | 1 | | | 11/23/03 | 29.6798 | -83.2567 |
| 179-03 | N. excitans | 1 | | | 11/25/03 | 28.7869 | -81.9818 |
| 099-04 | N. excitans | 1 | | | 07/14/04 | 31.4978 | -84.5933 |
| NS094 | N. excitans | 1 | | 2 | 10/18/14 | 29.7173 | -82.4570 |
| NS095 | N. excitans | 1 | | 1 | 10/18/14 | 29.7184 | -82.4562 |
| NS097 | N. excitans | 1 | | | 10/18/14 | 29.7351 | -82.4455 |
| NS098 | N. excitans | 1 | | | 10/18/14 | 29.7309 | -82.4520 |
| NS100.01 | N. excitans | | | 4 | 10/19/14 | 29.7074 | -82.4522 |
| NS100.02 | N. excitans | | | 5 | 10/19/14 | 29.7074 | -82.4522 |

| | | | | | | | |
|---|---|---|---|---|---|---|---|
| NS104 | N. excitans | | 7 | | 11/09/14 | 29.7076 | -82.4523 |
| NS108 | N. excitans | 1 | 9 | | 11/10/14 | 28.7870 | -81.9818 |
| NS133.02 | N. excitans | 1 | 4 | | 05/09/15 | 36.4248 | -77.6326 |
| NS134 | N. excitans | 1 | | | 05/09/15 | 36.4647 | -77.8229 |
| NS135.02 | N. excitans | 1 | 4 | | 05/09/15 | 36.4379 | -78.0875 |
| 109-04 | N. fabricii | 1 | | | 07/18/04 | 35.9332 | -86.5323 |
| 111-04B | N. fabricii | 1 | | | 07/19/04 | 36.1370 | -85.5031 |
| 115-04 | N. fabricii | 1 | | | 07/19/04 | 36.0222 | -85.0438 |
| NS023.02 | N. fabricii | | 1 | | 04/27/14 | 35.9292 | -86.5233 |
| NS128 | N. fabricii | 1 | | | 05/08/15 | 35.4745 | -80.2625 |
| NS133.01 | N. fabricii | 2 | | 2 | 05/09/15 | 36.4248 | -77.6226 |
| NS135.01 | N. fabricii | 1 | 2 | | 05/09/15 | 36.4378 | -78.0875 |
| NS135.03 | N. fabricii | 2 | 3 | | 05/09/15 | 36.4378 | -78.0875 |
| NS138.02 | N. fabricii | 1 | | | 05/10/15 | 37.2094 | -77.4743 |
| NS142 | N. fabricii | 1 | | | 05/25/15 | 38.4545 | -76.0661 |
| NS143 | N. fabricii | 2 | 4 | | 05/25/15 | 38.0627 | -76.5344 |
| NS144 | N. fabricii | 3 | 2 | | 05/25/15 | 37.5005 | -76.3014 |
| NS023.01 | N. hetricki | | 5 | | 04/27/14 | 35.9292 | -86.5233 |
| NS031 | N. hetricki | 2 | | | 05/05/14 | 37.6843 | -77.0825 |
| NS113.02 | N. hetricki | 3 | 8 | | 04/25/15 | 36.1397 | -85.8066 |
| NS114.02 | N. hetricki | 5 | 7 | | 04/25/15 | 35.8104 | -86.3984 |
| NS120.01 | N. hetricki | | 5 | | 05/09/15 | 36.5533 | -78.1801 |
| NS136.01 | N. hetricki | 1 | 3 | | 05/09/15 | 36.5533 | -78.1801 |
| NS136.02 | N. hetricki | 1 | 1 | | 05/09/15 | 36.5533 | -78.1801 |
| NS138.01 | N. hetricki | 1 | 3 | | 05/10/15 | 37.2094 | -77.4743 |
| NS138.02 | N. hetricki | 1 | 3 | | 05/10/15 | 37.2094 | -77.4743 |
| NS138.03 | N. hetricki | 1 | | | 05/10/15 | 37.2094 | -77.4743 |
| NS140.03 | N. hetricki | 2 | | | 05/10/15 | 37.1959 | -77.5244 |
| 082-04 | N. knereri | 1 | | | 07/11/04 | 29.0046 | -81.7574 |
| NS101 | N. knereri | 1 | | | 10/20/14 | 29.0046 | -81.7581 |
| NS102 | N. knereri | 1 | 3 | | 10/20/14 | 29.0340 | -81.6404 |
| NS214 | N. knereri | 2 | 5 | | 07/11/16 | 29.0040 | -81.7577 |
| NS217 | N. lecontei | 1 | | | 08/15/16 | 48.0302 | -84.6490 |
| 171-02 | N. lecontei | 1 | | | 08/10/02 | 44.7305 | -79.1686 |
| 174-02 | N. lecontei | 1 | | | 08/10/02 | 44.7305 | -79.1686 |
| 175-02 | N. lecontei | 1 | | | 08/10/02 | 44.7305 | -79.1686 |
| 345-02 | N. lecontei | 1 | | | 08/19/02 | 46.3949 | -79.2442 |
| 372-02 | N. lecontei | 1 | | | 09/25/02 | 41.8741 | -70.6524 |
| 075-04 | N. lecontei | 1 | | | 07/10/04 | 28.0955 | -81.2752 |
| 076-04 | N. lecontei | 1 | | | 07/10/04 | 28.0955 | -81.2752 |
| 087-04 | N. lecontei | 1 | | | 07/13/04 | 29.7476 | 82.4775 |
| 106-04 | N. lecontei | 1 | | | 07/17/04 | 32.0738 | -83.7606 |
| 188-04 | N. lecontei | 1 | | | 08/15/04 | 44.7589 | -91.4570 |

| | | | | | | | | |
|---|---|---|---|---|---|---|---|---|
| 207-04 | N. lecontei | 1 | | | | 08/17/04 | 45.9753 | -90.4964 |
| NS043.03 | N. lecontei | | | 8 | | 07/02/14 | 45.8223 | -91.8884 |
| NS050 | N. lecontei | 1 | | | | 07/04/14 | 44.8629 | -89.6366 |
| NL006 | N. lecontei | 1 | | | | 11/08/14 | 30.5931 | -84.3652 |
| NL007 | N. lecontei | 1 | | | | 11/08/14 | 30.2906 | -84.3514 |
| NS145.01 | N. lecontei | 2 | | | | 06/13/15 | 35.9803 | -85.0152 |
| NS145.03 | N. lecontei | 2 | | | | 06/13/15 | 35.9803 | -85.0152 |
| NS145.04 | N. lecontei | 1 | | | | 06/13/15 | 35.9803 | -85.0152 |
| NS145.05 | N. lecontei | | 1 | | | 06/13/15 | 35.9803 | -85.0152 |
| NS158 | N. lecontei | 2 | | | | 07/09/15 | 38.1003 | -83.5035 |
| NS160.01 | N. lecontei | 1 | | | | 07/23/15 | 40.3681 | -74.3026 |
| NS184.02 | N. lecontei | 1 | | | | 08/17/15 | 46.1149 | -90.5511 |
| NS186.01 | N. lecontei | 2 | | 4 | | 08/23/15 | 35.9803 | -85.0152 |
| NS186.02 | N. lecontei | 1 | | | | 08/23/15 | 35.9803 | -85.0152 |
| NS186.03 | N. lecontei | 1 | | | | 08/23/15 | 35.9803 | -85.0152 |
| NS204 | N. lecontei | 1 | | | | 06/28/16 | 45.8223 | -91.8884 |
| NS212.01 | N. lecontei | 5 | | 6 | | 07/10/16 | 29.0978 | -82.1861 |
| NS216 | N. lecontei | 3 | | 8 | | 07/11/16 | 29.3207 | -81.7267 |
| NS037 | N. maurus | 3 | | 7 | | 06/17/14 | 45.6643 | -89.4919 |
| NMr002 | N. maurus | 4 | | 6 | | 07/02/14 | 45.9975 | -91.6183 |
| NMr003 | N. maurus | 1 | | 1 | | 07/03/14 | 45.6019 | -89.3511 |
| NS203 | N. maurus | 3 | | | | 06/28/16 | 45.5792 | -91.759 |
| NS205.02 | N. maurus | 1 | | 3 | | 06/28/16 | 45.8428 | -91.8877 |
| NS206.02 | N. maurus | 3 | | | | 06/29/16 | 46.5544 | -91.3228 |
| NS208 | N. maurus | 1 | | 3 | | 06/29/16 | 46.1150 | -90.5513 |
| NS210.01 | N. maurus | 1 | | | | 06/29/16 | 46.0538 | -89.4857 |
| NS210.02 | N. maurus | 2 | | 2 | | 06/29/16 | 46.0538 | -89.4857 |
| NS105 | N. merkeli | 1 | | 3 | | 11/10/14 | 29.9238 | -81.5317 |
| NS106 | N. merkeli | 1 | | 2 | | 11/10/14 | 26.9260 | -81.5177 |
| NS107 | N. merkeli | | | 1 | | 11/10/14 | 26.9214 | -81.5086 |
| NS109 | N. merkeli | 1 | | 3 | | 11/10/14 | 28.7870 | -81.9818 |
| NS212.02 | N. merkeli | 1 | | | | 07/10/16 | 29.0978 | -82.1861 |
| NS212.03 | N. merkeli | 1 | | 1 | | 07/10/16 | 29.0978 | -82.1861 |
| NS213 | N. merkeli | 1 | | | | 07/10/16 | 28.7864 | -81.9816 |
| 214-02 | N. nigroscutum | 1 | | | | 08/13/02 | 48.3763 | -89.5844 |
| 209-04 | N. nigroscutum | 1 | | | | 08/17/04 | 46.0865 | -90.5507 |
| NS051 | N. nigroscutum | 1 | | 1 | | 07/05/14 | 44.8440 | -89.6906 |
| NS070.03 | N. nigroscutum | | | 2 | | 08/18/14 | 45.1248 | -92.3836 |
| NS205.01 | N. nigroscutum | 4 | | 1 | | 06/28/16 | 45.8428 | -91.8877 |
| NS206.01 | N. nigroscutum | 4 | | 1+1* | | 06/29/16 | 46.5544 | -91.3228 |
| NS211 | N. nigroscutum | 3 | 1 | 2+1* | | 06/30/16 | 44.8440 | -89.6905 |
| NP003 | N. pinetum | | | | 16 | 07/2313 | 36.0023 | -85.0438 |
| NP034 | N. pinetum | | | 2 | | 08/15/15 | 38.0324 | -84.5655 |

| | | | | | | | | |
|---|---|---|---|---|---|---|---|---|
| NP036 | N. pinetum | | | 2 | | 08/19/15 | 38.0324 | -84.5655 |
| NP039 | N. pinetum | | | 7 | | 08/20/15 | 37.9706 | -84.4976 |
| NP041 | N. pinetum | | | 1 | | 08/20/15 | 37.9729 | -84.5004 |
| 377-02 | N. pinetum | 1 | | | | 09/25/02 | 41.8459 | -70.6798 |
| 113-04 | N. pinetum | 1 | | | | 07/19/04 | 35.98 | -85.0152 |
| NP047 | N. pinetum | | | 6 | | 09/06/15 | 39.6188 | -83.6037 |
| NP048 | N. pinetum | 1 | | | | 09/06/15 | 39.6188 | -83.6037 |
| NP049 | N. pinetum | 1 | | | | 09/06/15 | 39.6188 | -83.6037 |
| NP050 | N. pinetum | 2 | | | | 09/06/15 | 39.6188 | -83.6037 |
| NP051 | N. pinetum | 1 | | | | 09/06/15 | 39.6188 | -83.6037 |
| NP052 | N. pinetum | 1 | | 4 | | 09/06/15 | 39.6188 | -83.6037 |
| NP053 | N. pinetum | 1 | | | | 09/06/15 | 39.6188 | -83.6037 |
| 373-02 | N. pinusrigidae | 1 | | | | 09/25/02 | 41.8459 | -70.6776 |
| 375-02 | N. pinusrigidae | 1 | | | | 09/25/02 | 41.8459 | -70.6776 |
| NS188.02 | N. pinusrigidae | 1 | 1 | 10 | | 09/13/15 | 39.8864 | -74.5099 |
| NS188.03 | N. pinusrigidae | 1 | 1 | | | 09/13/15 | 39.8864 | -74.5099 |
| NS188.04 | N. pinusrigidae | 1 | 1 | 12 | | 09/13/15 | 39.8864 | -74.5099 |
| NS189.01 | N. pinusrigidae | 1 | 1 | | | 09/13/15 | 39.8860 | -74.5061 |
| NS189.02 | N. pinusrigidae | 1 | 1 | 4 | | 09/13/15 | 39.8860 | -74.5061 |
| NS189.03 | N. pinusrigidae | 1 | 1 | 12 | | 09/13/15 | 39.8860 | -74.5061 |
| NS189.04 | N. pinusrigidae | 1 | 1 | 4 | | 09/13/15 | 39.8860 | -74.5061 |
| 076-02 | N. pratti | 1 | | | | 06/29/02 | 46.3113 | -81.6553 |
| 103-02 | N. pratti | 1 | | | | 06/30/02 | 45.9298 | -77.3318 |
| NS022 | N. pratti | 6 | | 6 | 3 | 04/13/14 | 38.0847 | -83.5098 |
| NS034 | N. pratti | 1 | | 4 | | 06/16/14 | 46.1113 | -89.6693 |
| NS110 | N. pratti | 2 | | | | 04/24/15 | 36.8544 | -84.8457 |
| NS112 | N. pratti | 1 | | 1 | | 04/24/15 | 36.5753 | -85.1380 |
| NS117.01 | N. pratti | 6 | | 6 | | 04/25/15 | 35.9419 | -86.5278 |
| NS118.02 | N. pratti | 1 | | 1 | | 04/25/15 | 35.9419 | -86.5278 |
| NS118.03 | N. pratti | 1 | | | | 04/25/15 | 35.9419 | -86.5278 |
| NS122 | N. pratti | 2 | | 1 | | 04/26/15 | 34.5275 | -84.9465 |
| NS123.01 | N. pratti | 3 | | | | 05/06/15 | 38.0860 | -83.5117 |
| NS123.02 | N. pratti | 1 | | | | 05/06/15 | 38.0860 | -83.5117 |
| NS123.03 | N. pratti | 3 | | 4 | | 05/06/15 | 38.0860 | -83.5117 |
| NS124 | N. pratti | | | 4 | | 05/06/15 | 38.0860 | -83.5117 |
| NS125 | N. pratti | | | 4 | | 05/06/15 | 38.0843 | -83.5098 |
| NS140.01 | N. pratti | 1 | | 1 | | 05/06/15 | 37.1959 | -77.5244 |
| NS138.04 | N. pratti | 1 | | | | 05/10/15 | 37.2094 | -77.4743 |
| NS150 | N. pratti | 1 | | | | 06/14/15 | 47.9385 | -83.0621 |
| NS153 | N. pratti | 1 | | | | 06/27/15 | 46.6237 | -81.4547 |
| NS154.02 | N. pratti | 4 | | 4 | | 06/27/15 | 46.3111 | -81.6557 |
| NS155 | N. pratti | 2 | | 1 | | 06/28/15 | 46.3023 | -79.3835 |
| NS163 | N. pratti | 2 | | 6 | | 07/25/15 | 44.4801 | -67.5931 |

| | | | | | | | |
|---|---|---|---|---|---|---|---|
| NS164 | N. pratti | 1 | | | 07/25/15 | 44.7799 | -67.5930 |
| NS165 | N. pratti | 1 | | 3 | 07/25/15 | 44.4774 | -67.5908 |
| NS166 | N. pratti | 1 | | | 07/25/15 | 44.4775 | -67.5906 |
| NS207 | N. pratti | 2 | | | 06/29/16 | 46.5086 | -90.5015 |
| 155-02 | N. rugifrons | 1 | | | 08/08/02 | 45.9316 | -77.3332 |
| 156-02 | N. rugifrons | 1 | | | 08/08/02 | 45.9316 | -77.3332 |
| 157-02 | N. rugifrons | 1 | | | 08/08/02 | 45.9316 | -77.3332 |
| 181-02 | N. rugifrons | 1 | | | 08/10/02 | 45.7987 | -80.5352 |
| 182-02 | N. rugifrons | 1 | | | 08/10/02 | 45.7987 | -80.5352 |
| 183-02 | N. rugifrons | 1 | | | 08/10/02 | 45.7987 | -80.5352 |
| 184-02 | N. rugifrons | 1 | | | 08/10/02 | 45.7987 | -80.5352 |
| 190-02 | N. rugifrons | 1 | | | 08/11/02 | 45.9718 | -80.5760 |
| 204-02 | N. rugifrons | 1 | | | 08/11/02 | 46.6240 | -81.4544 |
| 208-02 | N. rugifrons | 1 | | | 08/11/02 | 46.6240 | -81.4544 |
| 242-02 | N. rugifrons | 1 | | | 08/14/02 | 49.7247 | -94.2481 |
| 243-02 | N. rugifrons | 1 | | | 08/14/02 | 49.7247 | -94.2481 |
| 267-02 | N. rugifrons | 1 | | | 08/16/02 | 48.0295 | -84.6518 |
| 268-02 | N. rugifrons | 1 | | | 08/16/02 | 48.0295 | -84.6518 |
| 273-02 | N. rugifrons | 1 | | | 08/16/02 | 48.0458 | -84.5503 |
| 274-02 | N. rugifrons | 1 | | | 08/16/02 | 48.0458 | -84.5503 |
| 277-02 | N. rugifrons | 1 | | | 08/16/02 | 47.9763 | -84.5318 |
| 283-02 | N. rugifrons | 1 | | | 08/17/02 | 48.0934 | -82.6727 |
| 287-02 | N. rugifrons | 1 | | | 08/17/02 | 48.1851 | -82.2276 |
| 292-02 | N. rugifrons | 1 | | | 08/17/02 | 48.2203 | -82.0736 |
| 298-02 | N. rugifrons | 1 | | | 08/17/02 | 48.3651 | -81.5758 |
| 299-02 | N. rugifrons | 1 | | | 08/17/02 | 47.8084 | -81.5917 |
| 304-02 | N. rugifrons | 1 | | | 08/17/02 | 47.8084 | -81.5917 |
| 310-02 | N. rugifrons | 1 | | | 08/18/02 | 47.4835 | -81.8459 |
| 313-02 | N. rugifrons | 1 | | | 08/18/02 | 47.4835 | -81.8459 |
| 314-02 | N. rugifrons | 1 | | | 08/18/02 | 47.4771 | -81.5260 |
| 318-02 | N. rugifrons | 1 | | | 08/18/02 | 47.4771 | -81.5260 |
| 319-02 | N. rugifrons | 1 | | | 08/18/02 | 47.4849 | -81.4034 |
| 324-02 | N. rugifrons | 1 | | | 08/18/02 | 47.4849 | -81.4034 |
| 333-02 | N. rugifrons | 1 | | | 08/18/02 | 47.6651 | -80.6184 |
| NS044 | N. rugifrons | | | 6 | 07/02/14 | 45.8396 | -91.9139 |
| NS052 | N. rugifrons | 1 | | 4 | 07/20/14 | 45.8396 | -91.9139 |
| NS070.01 | N. rugifrons | | | 1 | 08/18/14 | 45.1248 | -92.3836 |
| NS172.04 | N. rugifrons | | | 3 | 08/14/15 | 47.4648 | -81.8467 |
| NS200 | N. rugifrons | 1 | 1 | 2 | 06/28/16 | 44.5711 | -91.6363 |
| 193-02 | N. swainei | 1 | | | 08/11/02 | 46.0580 | -80.6252 |
| 195-02 | N. swainei | 1 | | | 08/11/02 | 46.1334 | -80.6873 |
| 254-02 | N. swainei | 1 | | | 08/15/02 | 48.6885 | -93.0019 |
| 301-02 | N. swainei | 1 | | | 08/17/02 | 47.8084 | -81.5917 |

| | | | | | | | |
|---|---|---|---|---|---|---|---|
| 307-02 | N. swainei | 1 | | | 08/17/02 | 47.8084 | -81.5917 |
| 336-02 | N. swainei | 1 | | | 08/18/02 | 47.6387 | -79.9771 |
| 337-02 | N. swainei | 1 | | | 08/18/02 | 47.6387 | -79.9771 |
| 179-04 | N. swainei | 1 | | | 08/14/04 | 44.5814 | -91.5290 |
| 185-04 | N. swainei | 1 | | | 08/14/04 | 44.5709 | -91.6352 |
| 211-04b | N. swainei | 1 | | | 08/17/04 | 46.1195 | -90.5544 |
| NS053.01 | N. swainei | | 5 | | 07/20/13 | 46.1143 | -90.5513 |
| NS056.02 | N. swainei | | 1 | | 07/20/14 | 44.8440 | -89.6906 |
| NS061.02 | N. swainei | | 5 | | 08/04/14 | 47.6537 | -81.0517 |
| NS091 | N. swainei | 1 | | | 09/13/14 | 45.9974 | -91.6181 |
| NS169.02 | N. swainei | 1 | | | 08/13/15 | 45.9315 | -77.3333 |
| NS184.03 | N. swainei | 3 | | | 08/17/15 | 46.1149 | -90.5511 |
| NS184.04 | N. swainei | 1 | | | 08/17/15 | 46.1149 | -90.5511 |
| NTL003 | N. taedae | | 1 | | 05/23/14 | 36.0095 | -87.3781 |
| NTL001 | N. taedae | | 2 | | 05/18/14 | 36.3177 | -94.7105 |
| NTL004 | N. taedae | | 4 | | 06/09/14 | 37.7271 | -92.7033 |
| NS113.01 | N. taedae | 3 | | | 04/25/15 | 36.1397 | -85.8066 |
| NS114.01 | N. taedae | 8 | 3 | | 04/25/15 | 36.1791 | -85.9456 |
| NS115.03 | N. taedae | | 5 | | 04/25/15 | 36.4716 | -86.6816 |
| NS117.02 | N. taedae | 5 | 8 | | 04/25/15 | 35.9419 | -86.5278 |
| NS118.01 | N. taedae | 1 | | | 04/25/15 | 35.9419 | -86.5278 |
| NS119 | N. taedae | 2 | | | 04/25/15 | 35.9333 | -86.5322 |
| LL185 | N. virginiana | | 1 | | 07/12/15 | 37.3585 | -77.9611 |
| LL186 | N. virginiana | | 4 | | 07/12/15 | 37.3586 | -77.9609 |
| LL187 | N. virginiana | | 3 | | 07/12/15 | 37.3586 | -77.9609 |
| LL188 | N. virginiana | | 3 | | 07/12/15 | 37.3586 | -77.9609 |
| LL189 | N. virginiana | | 5 | | 07/12/15 | 37.3585 | -77.9611 |
| LL192 | N. virginiana | | 3 | | 07/12/15 | 37.9863 | -84.4171 |
| NS159 | N. virginiana | 1 | | 2 | 07/22/15 | 39.7042 | -78.3288 |
| NS099 | N. warreni | 1 | 3 | | 10/19/14 | 29.7093 | -82.4542 |
| NS103 | N. warreni | 1 | 3 | | 11/09/14 | 29.7091 | -82.4541 |

* Indicate videos using larvae from multiple collection sites; details can be found in Supplementary Table 4

Supplemental Table 4. Summary of videos using larvae from multiple collection sites.

| Video Name | Species | Colonies Used |
|---|---|---|
| NS086_088-1 | *N. abbotii* | NS086, NS088 |
| CN001 | *N. compar* | NS174, NS182 |
| CN002 | *N. compar* | NS175, NS184 |
| CN003 | *N. compar* | NS168, NS176, NS178 |
| CN004 | *N. compar* | NS169, NS170, NS172, NS174 |
| CN006 | *N. compar* | NS218, NS220, NS221 |
